# Decoding of the other’s focus of attention by a temporal cortex module

**DOI:** 10.1101/681957

**Authors:** H. Ramezanpour, P. Thier

## Abstract

Faces attract the observer’s attention towards objects and locations of interest for the other, thereby allowing the two agents to establish joint attention. Previous work has delineated a network of cortical “patches” in the macaque cortex, processing faces, eventually also extracting information on the other’s gaze direction. Yet, the neural mechanism that links information on gaze direction, guiding the observer’s attention to the relevant object has remained elusive. Here we present electrophysiological evidence for the existence of a distinct “gaze-following patch (GFP)” with neurons that establish this linkage in a highly flexible manner. The other’s gaze and the object, singled out by the gaze, are linked only if this linkage is pertinent within the prevailing social context. The properties of these neurons establish the GFP as a key switch in controlling social interactions based on the other’s gaze.

**One Sentence Summary:** Neurons in a “gaze-following patch” in the posterior temporal cortex orchestrate the flexible linkage between the other’s gaze and objects of interest to both, the other and the observer.

## Main Text

We use the other’s gaze direction to shift attention to the object the other one is attending to, thereby establishing joint attention. Joint attention allows us to develop a theory of (the other’s) mind (ToM) (*1*) by mapping one’s own thoughts, beliefs and desires associated with the attended object onto the other one. Although it is questionable if monkeys also dispose of a full ToM, they follow the other’s gaze to establish joint attention (*2–4*). An important distinction between human and non-human primate gaze-following is the different weight of eye and head gaze cues. Not surprisingly in view of the fact that the eyes of non-human primates lack the conspicuous features of the human eye (*5*), monkeys’ gaze-following relies primarily on head gaze rather than on eye gaze cues (*4*). This important difference notwithstanding, the evidence available emphasizes close similarities of human and non-human gaze-following behavior, suggesting the possibility of a homologous system shared within the primate order. For instance, comparative fMRI work has delineated a distinct cortical node in the posterior temporal cortex of both rhesus monkeys and man specifically activated by gaze-following. In both species this *gaze-following patch* (GFP) is located in the immediate vicinity of the more posterior elements of the so-called face patch system (*6, 7*), a system that has been implicated in the extraction of different aspects of information on faces such as identity, face or head orientation or facial expression (*8–11*). Actually, not only face orientation is important for the guidance of the observer’s gaze but also information on identity or facial expressions as both are known to modulate human gaze-following (*12*). Hence, it is likely that the GFP may draw on information from the face patch system. Yet, the neural mechanisms that may allow the GFP to use information on the other’s facial features into a gaze-following response establishing joint attention are not known. It is also unclear if neurons in the GFP may possibly contribute to the cognitive control of gaze-following, integrating contextual information relevant for the modulation of the behavior. After all, although human and monkey gaze-following has features of a quasi reflex-like behavior that kicks in at short latency, it can be suppressed to a considerable degree if not appropriate within a given context (*13, 14*).

With these questions in mind, we explored the GFP of rhesus monkeys and adjoining regions of the superior temporal cortex (STS), deploying tasks that asked the observer to follow the other’s gaze or to suppress gaze-following if not expedient. Our results suggest that neurons in the GFP link information on the other’s gaze and the object singled out by the gaze, provided that this linkage is pertinent within the prevailing social context.

## Results

We recorded the activity of well-isolated single neurons in the GFP and adjacent regions of the right STS of two rhesus monkeys. In one of the two, the location of the GFP had been delineated in preceding fMRI experiments in which we had searched for BOLD activity associated with gaze-following as described by (*6*). In the second monkey, we relied on the same coordinates as reference, when exploring the STS. Both monkeys had learned to follow the direction of a monkey head (“demonstrator”) presented on a monitor. The demonstrator turned to one out of four spatial targets ((head) gaze-following task). Alternatively, the monkeys had to use the facial identity of the portrayed monkeys to determine the relevant target. To this end, they had to rely on a learned association between the four targets and the four possible identities (identity-mapping task). An instructive color cue presented on a baseline portrait before the appearance of the 4 spatial cues and targets told the monkey whether to deploy the gaze or the identity rule when dealing with the monkey portraits. The two trial types were presented randomly interleaved (**Fig. 1A, B**). The design of the paradigm allowed us to dissociate neural activity evoked by features of the portraits from activity associated with the shift of attention to a particular target object, prompted by two different social cues, gaze direction and facial identity respectively. As the portraits and the overt behavior they caused were the same, independent of the rule, any difference in neural responses had to be a reflection of differences between the cognitive processes responsible for the rule-based selection of target objects. Monkey L reached a mean performance of 84 ± 5% correct trials on the gaze-following task and 82 ± 7% on the identity-mapping task, whereas monkey T attained 71 ± 8% and 73± 11% respectively (mean ± std) (**Fig. 1c**). Both monkeys’ performance was significantly above the 25% chance level (P < 0.001, binomial test) and independent of the specific task (Wilcoxon signed-rank test, p> 0.05). We also tested the responses of the same neurons to the passive viewing of faces and a variety of biological and non-biological objects (**Fig. 1D**).

**Fig. 1.**
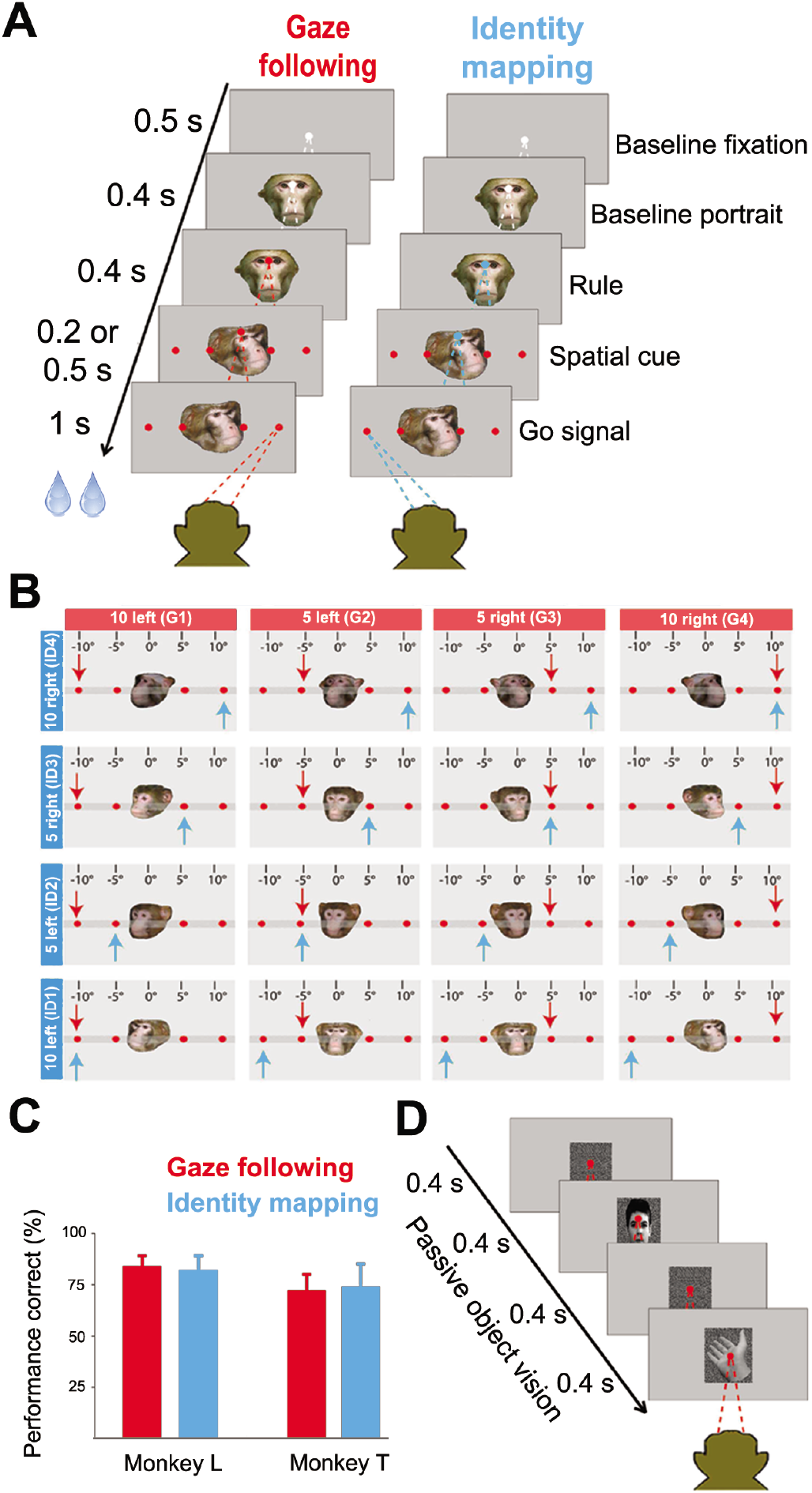
Behavioral Paradigms. (**A**) Each trial started with the presentation of a white central fixation dot, 500ms later supplemented by a straight ahead looking portrait. 400ms after the onset of the portrait, the fixation dot changed color, specifying the rule to be applied on the trial. Red indicated gaze-following, green identity-mapping (note: rendered blue in all figures for better visibility). 400ms later, the straight ahead portrait was replaced by the portrait of another monkey (“demonstrator”) looking at one out of the four targets, looming up at the same time. The disappearance of the central fixation point, 200 or 500 ms later – depending on the day - served as the go signal to make a saccade to the target specified by the demonstrator. In the gaze-following task the relevant cue was the demonstrator’s head gaze whereas in the identity-mapping task, the observer was asked to ignore the gaze direction and to make a saccade and respond to the target specified by the portrait’s identity, resorting to learned associations between the 4 targets and the 4 possible identities of the demonstrators. These two tasks were randomly interleaved. (**B**) The 4×4 cue matrix defined by the 4 possible demonstrator identities and the 4 possible orientations of the demonstrator’s head (40° left, 20° left, 10° right, 20° right from the demonstrator’s viewpoint corresponding to targets at 10° left, 5° left, 5° right, 10° right from the perspective of the observer). The blue arrow in each cell specifies the target to be chosen according to the prevailing identity, the red arrow the one singled out by head gaze. (**C**) The behavioral performance of the two monkeys in each of the two behavioral paradigms was very good, well above the 25% chance level, not significantly different for the two tasks (p>0.05) and without significant difference between the two monkeys. Error bars represent standard error. (**D**) The passive viewing task required the observer to fixate a 0.2 ° dot while exposed to a sequence of images of faces and non-face stimuli, presented randomly interleaved. Each image was on for 400 ms and followed by a 400 ms duration random dot background.

### Single pSTS neurons encode gazed-at targets

Altogether, we tested 923 neurons recorded from the posterior STS (pSTS) of the two monkeys on all three tasks. Out of these, 426 neurons (172 neurons in monkey T and 254 neurons in monkey L) exhibited significant changes of their discharge rate relative to baseline (Kruskall-Wallis ANOVA, p<0.05) in at least one of two key phases of a trial, namely during the presentation of the rule and/ or during the subsequent availability of the spatial information provided by gaze direction or facial identity. In total, 109 (out of 426 task-related) neurons exhibited selectivity for head gaze (“gaze-following (GF) neurons”), 37 neurons for facial identity specifying spatial locations (“identity-mapping (IM) neurons”) and 12 neurons were responsive to both gaze and identity (“mixed selectivity neurons”)(see pie chart in **Fig. 2A**). **Fig. 2B** depicts the distribution of spatial preferences of GF and IM neurons based on the target yielding the maximal response. It shows that all four possible targets are well represented in the data set without any bias for the left or the right side. The discharge profiles of two exemplary spatially-selective neurons and one exemplary classical face-selective neuron lacking interest in spatial information are assembled in **Fig. 2C-E. Fig. 2C** shows a typical GF neuron. Its discharge profile was characterized by very similar discharge rates in the gaze following and the identity mapping tasks until the time the monkey portrait provided information on the target location to be chosen. In case the rule demanded gaze-following, the discharge rate was significantly higher than for identity-mapping if the cued target was the one at 10° on the right (target G4, corresponding to 40° left from the view of the demonstrator monkey). The difference became significant shortly after the onset of the spatial cue, reached its maximum 145 ms later (first peak) and stayed until the time of the indicative saccade. **Fig. 2D** depicts another neuron exhibiting a qualitatively similar discharge pattern, yet with some preference for identity-mapping defining the target at 10° on the left (target ID1). Both neurons lacked specificity for faces when tested for visual responses to the presentation of faces and a variety of biological and non-biological objects during stationary fixation (“object vision task”). On the other hand, the neuron shown in **Fig. 2E** was a classical face-selective neuron when tested in the object vision task, characterized by a strong preference for face stimuli. Clear bursts of activity evoked by the appearance of the portraits also characterized the active tasks without any difference between the two conditions or between the spatial targets within each task.

**Fig. 2.**
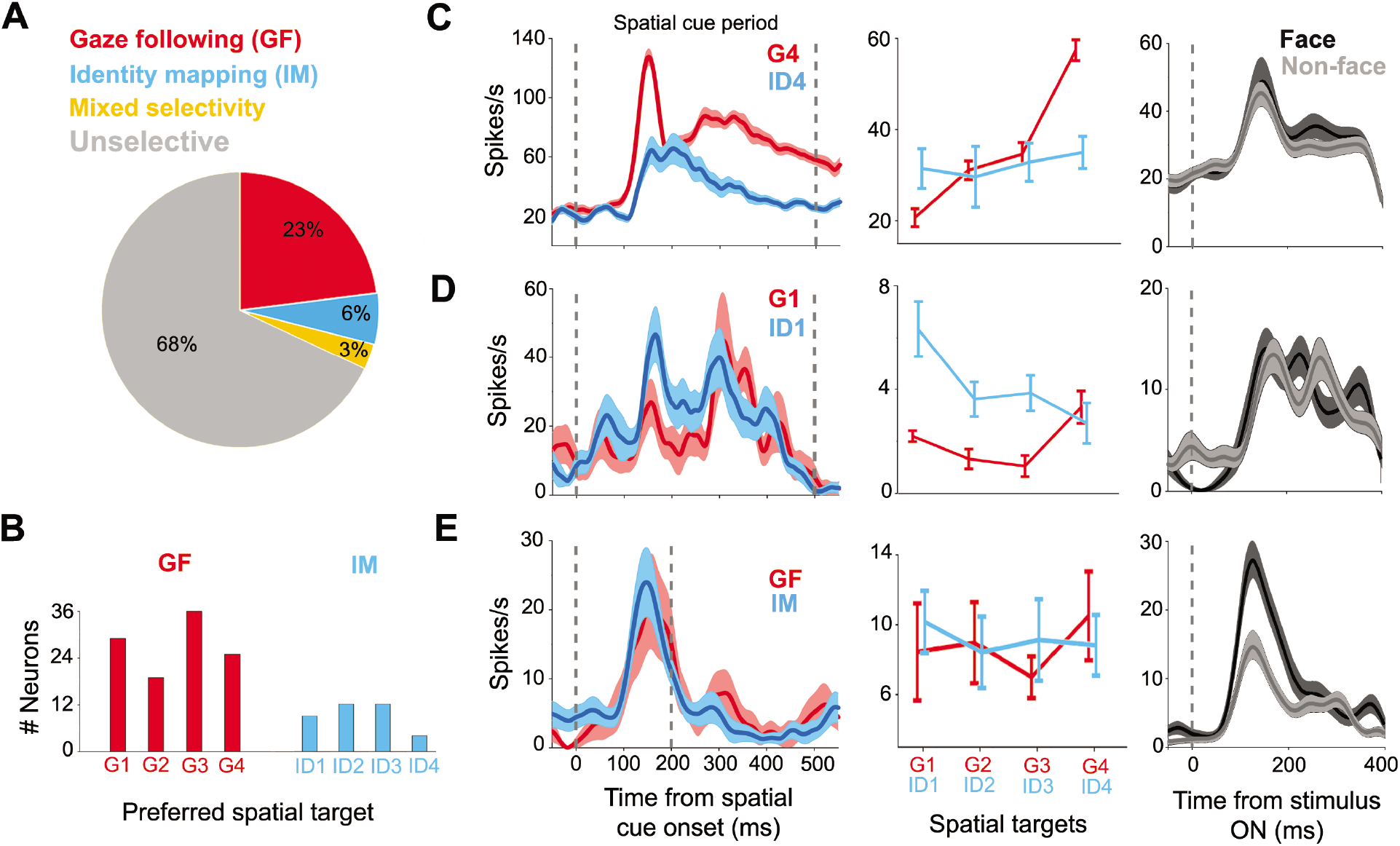
The variety of neuronal response features in the pSTS. (**A**) Breakdown of response preferences of neurons in the GFP during the spatial cueing period. (**B**) Distribution of targets preferred by spatially-selective GF and IM cells respectively, (G1=10° left, G2=5° left, G3= 5° right, G4=10° right, ID1=10° left, ID2=5° left, ID3= 5° right, ID4=10° right) according to their most preferred targets pooled over both monkeys. (**C**) Response profiles of an exemplary spatially-selective GF neuron tested in the GF and the IM task (left and middle panel) and the passive viewing task (right panel). This neuron was activated in both the GF and the IM task, yet clearly more in the former with preference for target G4. It did not show face selectivity in the passive viewing task. (**D**) Exemplary IM neuron with preference for target ID1 in the identity-mapping task. Also this neuron failed to exhibit face selectivity in the passive viewing task. (**E**) Example of a classical face-selective neuron that preferred faces over non-face stimuli in the passive viewing task and a clear face response in both the gaze-following and the identity-mapping task without exhibiting any sensitivity to the other aspects of the two tasks. In all panels the vertical line at t=0 identifies the onset of the 4 targets while the second vertical line at 500 ms (C, D) or 200ms (E) identifies the time of the go signal. Error bars and shaded areas represent standard error in all figure parts.

A clear preference for distinct targets was also exhibited by the population tuning curve based on all 109 spatially-selective GF neurons. To assess the selectivity of the population for distinct spatial targets we ranked the strength of the responses of all individual GF neurons to the four gaze targets and calculated population responses for each rank. Rank 1 stood for the most preferred gaze target (highest mean discharge in the period of 50 ms after the onset of the spatial cues until the appearance of the go cue) and rank 4 for the least preferred gaze target. As can be seen in **Fig. 3A**, the rank-specific population responses were very distinct with the largest burst of activity for rank 1, a smaller one for rank 2 and clear activity suppression for the two lowest ranked targets. **Fig. 3B** compares the population responses of the GF neurons for the highest and lowest ranked targets in the GF task with the responses evoked by the same targets cued by facial identity. In case of identity-mapping, the difference between the population responses for the two targets associated with the most and the least preferred target in the gaze-following task was dramatically reduced to a non-significant level (Mann-Whitney U-test, p=0.32). Both discharge profiles, in each case averaging over all 4 identities, lay in between the rank 1 and 4 responses evoked by gaze cueing. The residual response modulation in the spatial cueing period, uninfluenced by target position, may reflect the need to process facial identity in this task. An analogous analysis for the IM neurons did not reveal any significant difference between the population responses to the rank 1 and rank 4 targets (Mann-Whitney U test, p=0.1)(**Fig. S1B**). In other words, at the population level IM neurons do not convey information on spatial targets. This may suggest that the significant preferences for particular identity-targets association, exhibited by 37 out of the 76 IM neurons may actually reflect identity tuning rather than spatial tuning.

**Fig. 3.**
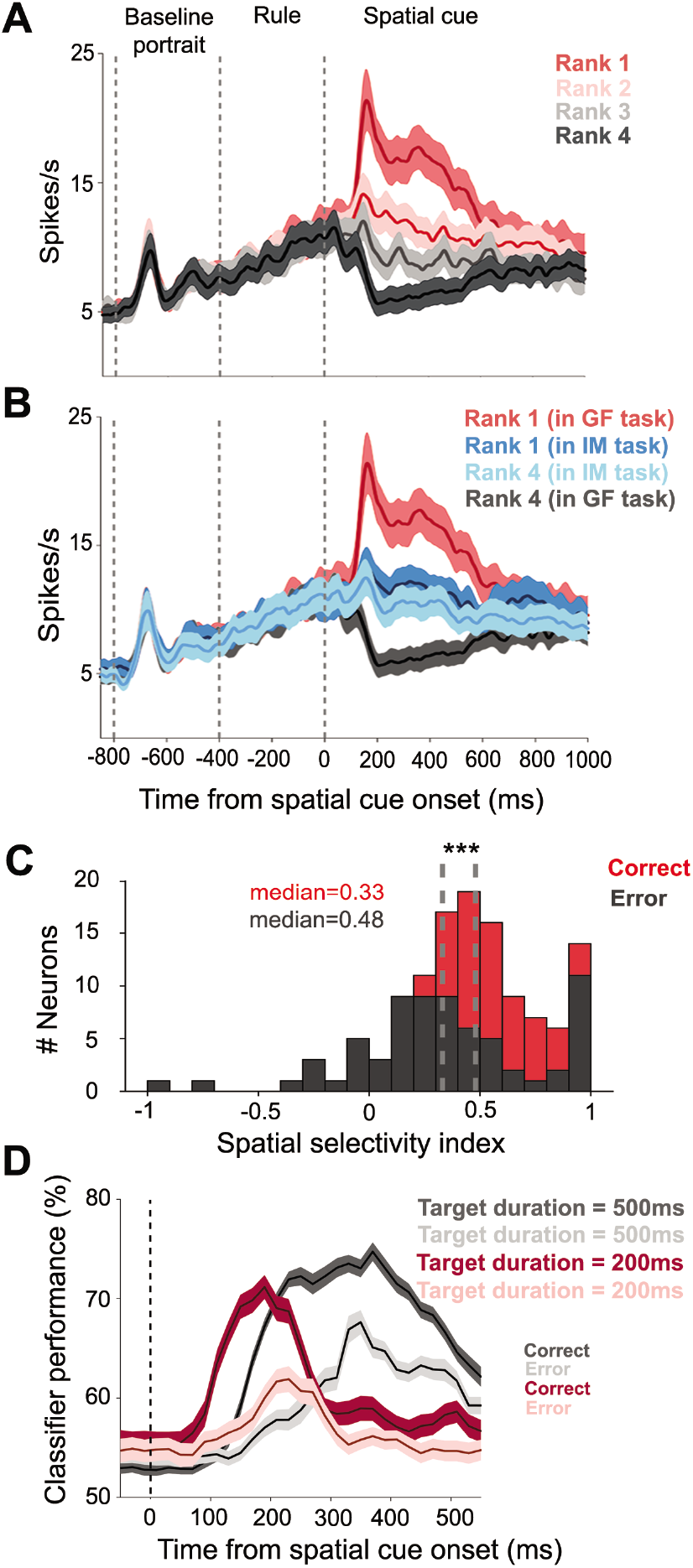
Population responses of spatially-selective GF neurons. (**A**) Population responses of 109 GF neurons from both monkeys for the target eliciting the strongest response (rank 1) in the spatial cue period, the second strongest (rank 2), the third (rank 3) and the fourth strongest response (rank 4). The population discharge associated with the two most preferred targets exhibited an increase in discharge rate, the ones associated with the least preferred targets a suppression. (**B**) Population responses of GF neurons for the most preferred and the least preferred targets in the GF task compared with the population responses to the same targets cued by identity in the IM task. (**C**) For the population of spatially-selective GF neurons, the median SSI values dropped significantly (Mann Whitney U test, p<0.001) from 0.48 for correct trials to 0.33 for error trials. SSI values for error trials could even become negative, indicating a reversal of preference, i.e. a target that was the most preferred one in the GF task became less effective than the least preferred one when studied in the IM task. (**D**) SVM decoding of the discrimination between the most preferred and least preferred target based on the activity of all spatially-selective GF, shown separately for correct and error trials and separately for the two durations of the spatial cueing periods (200 vs. 500 ms). Standard errors were obtained by deploying a bootstrapping procedure (n=1000). Shaded areas represent standard errors in all figures parts.

Under the assumption that the GF neurons underlie a monkey’s ability to follow gaze to the relevant target, error trials, in which the monkey fails to hit the target identified by the other’s gaze should be associated with reduced selectivity of the GF population discharge. To test this prediction, we calculated a spatial selectivity index (SSI), capturing the difference between the population responses to the most (rank 1) and the least preferred target (rank 4) divided by the sum. For the population of GF neurons, the distribution of SSI varied between 0 and 1 with a median of 0.48 for correct trials. For incorrect trials the whole distribution shifted to the left with a median of 0.33, significantly smaller than the one for correct trials (p < 0.001, Mann-Whitney U test; n = 109). Unlike the distribution for correct trials, the one for error trials spread into the negative range, indicating that quite a few neurons changed their spatial preferences (**Fig. 3C**). The notion that errors in gaze-following trials are a consequence of compromised selectivity of the GF neuron population signal is also supported by a time-resolved decoding analysis based on a linear support vector machine (SVM) classifier with 5-fold cross-validation, determining the amount of information available on the correct target. We obtained a decoding time course by performing this analysis in a 100 ms window and advancing in steps of 20 ms during the whole spatial cueing period. We performed this analysis separately for the two pools of GF neurons, tested with spatial cue windows of 200 ms (n=28) and 500 ms (n=81) respectively. As shown in **Fig. 3D**, information about the position of the spatial targets is present almost throughout the whole time. These results demonstrate that the population of GF neurons offers reliable information on the gazed-at target throughout a period from the onset of the spatial cue until the time of the go signal. For error trials, the maximum of the decoder classification performance dropped by about 10%, in line with the notion that precise shifts of attention to the gazed at target require a specific population signal.

To test if task-related neurons in the pSTS are indeed tuned only to social cues such as information on gaze direction or facial identity, we ran a control task with abstract symbols replacing the faces. Four specific symbols - a square, a circle, a triangle and a star had been learned to be associated with one out of the four possible targets each. 131 out of the pool of 426 GF or IM task-related neurons were tested on this task. Only very few (n=8) showed weak, albeit significant responses to targets cued by symbols. Moreover, the population response failed to distinguish correct and error trials. The same holds for a topographically distinct separate population of face-selective neurons (n=23). These neurons (with the exception of n=6) lacked spatial tuning in the gaze-following and identity-mapping tasks and as a group failed to discriminate between the correct and error trials (see Supplementary materials, **Fig. S1A**).

### pSTS neurons encode abstract rules and bias monkeys’ social choices

Spatial selectivity was not the only feature characterizing neurons in the pSTS. We could also identify 104 rule-selective neurons, either encoding the rule to follow gaze or to map identity. The population of rule-selective neurons overlapped with the one exhibiting spatial selectivity with 22% of the latter (43 out of 195) showing both spatial and rule selectivity. **Fig. 4A** depicts two exemplary rule-selective neurons, one preferring the gaze-following rule, the other one the identity-mapping rule. Both exhibited a clear increase of their discharge rates for the respective preferred rule, 131 and 159 ms (latency of first peak) respectively after the onset of the information on the prevailing rule. As exemplified by these two neurons, the rule-associated activation depended on whether the monkey was able to convert the rule into successful shifts of attention to the correct target or not. In case of error trials, the differential response of the GF neuron dropped dramatically, while, conversely, the IM neuron showed a higher amount of differentiation between the two rules. Note that both neurons lacked a significant response to the portraits comprising the non-preferred rule, i.e. similar to many spatially-selective neurons they were not sensitive to the vision of faces. The pie chart in **Fig. 4B** gives a breakdown of the numbers of rule-selective neurons in each category and each monkey. **Figure 4C** depicts the population responses of rule-selective neurons preferring the GF and the IM rule respectively. In both cases, the population plots exhibit excitatory responses to the preferred rule. However, whereas the population responses for neurons preferring the gaze-following rule show a weak excitatory response to the nonpreferred rule, the one for those preferring the identity-mapping rule is characterized by a discharge suppression in the presence of the non-preferred rule. Unlike identity-mapping, gaze-following has all the features of a domain specific cognitive function, among others characterized by a significant degree of automaticity. In other words, reliably suppressing the gaze-following reflex may need extra efforts involving the active suppression of the distracting gaze-following rule related activity.

**Fig. 4.**
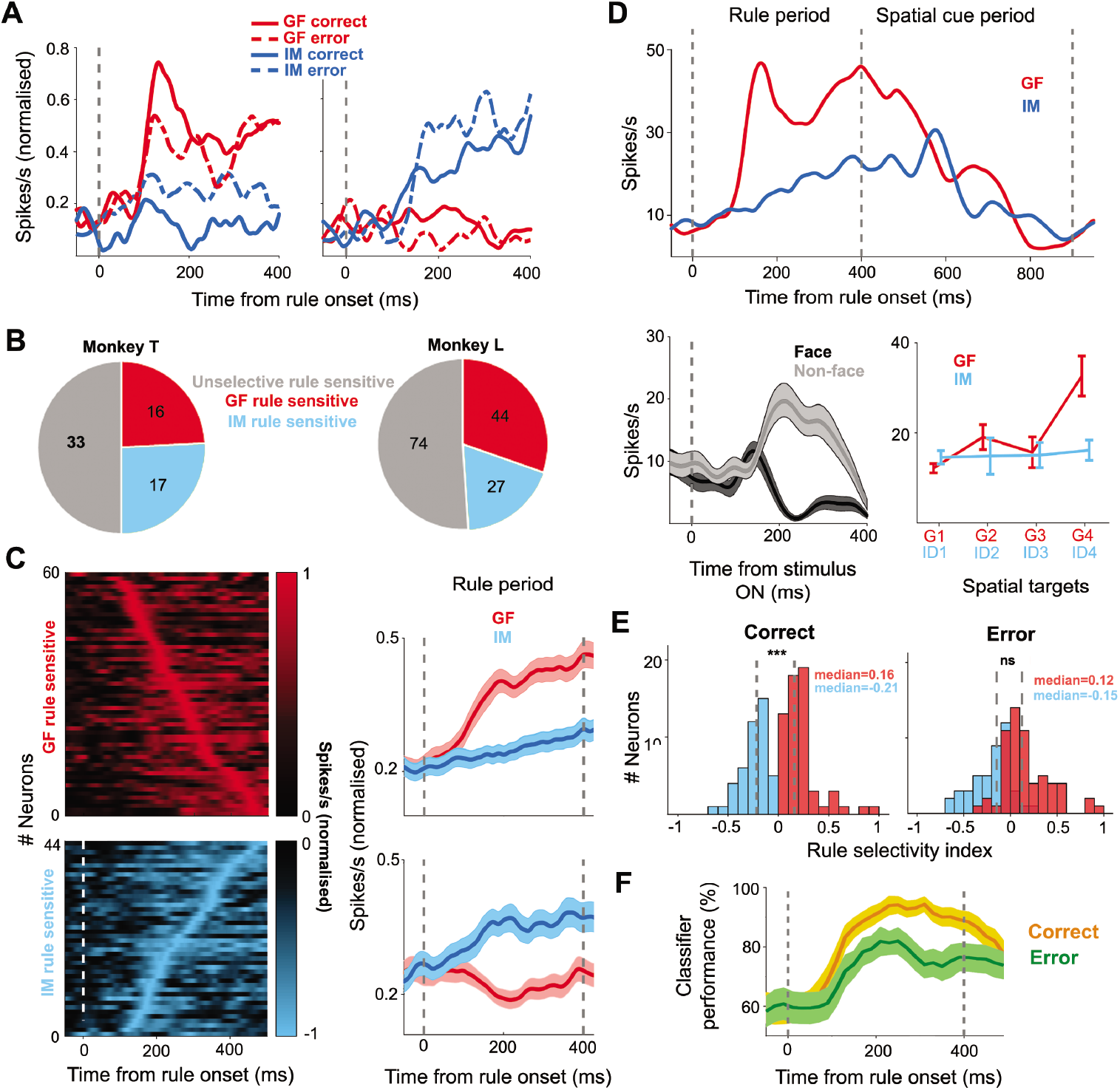
Rule selectivity of GFP neurons. (**A**) Responses of two exemplary GFP neurons to the presentation of the two rules, shown separately for correct and for error trials. The one on the left exhibited selectivity for the GF rule, while the one on the right preferred the IM rule. (**B**) Breakdown of rule preferences of rule sensitive GFP neurons for both monkeys. (**C**) The right panel depicts the time course of the population responses of the two pools of neurons preferring the GF rule (n=60) and the IM rule (n=44) respectively, tested in both the GF and the IM task. The panel on the left plots the contributions of individual neurons ordered according to the latency of their peak discharge rates. (**D**) Exemplary GFP neuron demonstrating that preferences for rules and spatial cues are yoked. This neuron preferred the gaze rule and exhibited a much stronger response to the gaze cue in the subsequent 500 ms of the spatial cueing period. The vertical line at 400ms denotes the onset of spatial cue. This neuron preferred non-face objects over faces in the passive viewing task. (**E**) Deterioration of the rule selectivity of the GFP as captured by the RSI results in erroneous decisions. (**F**) SVM decoding accuracy obtained from GFP rule-selective neurons for correct trails as compared to error trails. The shaded area represents standard deviations obtained by bootstrapping (n=1000). Note the clear drop in performance for error trials. Error bars and shaded areas represent standard error in figure parts except for (F) which shows standard deviations.

As alluded to above quite a few rule-selective neurons integrated rule selectivity and sensitivity to spatial targets, either identified by gaze or by identity, in any case in a congruent manner. That is to say that neurons that preferred the gaze rule also preferred gaze-following and vice versa neurons selective for the identity-mapping rule attentional shifts guided by identity. An example of a neuron selective for the gaze rule and for a target selected by gaze is depicted in **Fig. 4D**. We wondered if the degree of rule selectivity might predict the later spatial selectivity. This was indeed the case as shown by a quantitative analysis of the rule selectivity based on a rule selectivity index (RSI) calculated by the normalized difference of the mean discharges for the GF and IM conditions in the 400 ms of the rule window (details in the supplementary materials). In other words, the ability to decode the rule is relevant for the ability to shift attention to the right target. This is also indicated by a consideration of error trials. In the case of GF rule-selective neurons, median RSI values dropped from a median of 0.16 for correct trials to 0.12 for error trials (p < 0.05, Mann Whitney U test; n = 104)(**Fig. 4E**). Similarly, for the population of IM rule-selective neurons the median RSI values decreased from −0.21 for correct trials to −0.15 for error trials (p < 0.001, Mann Whitney U test; n = 104). As a consequence, although still significantly different ((p < 0.001, Mann Whitney U test, two-tailed; n = 104)) the distribution of RSI values for the two groups of neurons exhibited considerable overlap for error trials, an impairment that is in principle in accordance with the drop in behavioral selectivity (**Fig. 4E**). The same conclusion can be drawn from a decoding analysis deploying a support vector machine classifier. Here we asked how well the responses evoked by the two rules predicted the behavioral decisions. As shown in **Fig. 4F**, the classifier performance dropped significantly for erroneous decisions.

### Topography of the neural response types in the STS

We reconstructed the locations of recorded neurons in stereotactic coordinates based on a 3D rendering of the pSTS using anatomical MRI data sets available for two monkeys. The positions of neurons were then used to construct 2D density maps of response features following unfolding of the pSTS. To this end, we counted the number of neurons in each 0.5 mm^2^ of unfolded cortex and finally passed the resulting distribution through a 2D spatial filter (Gaussian, σ=2). **Fig. 5** depicts the resulting density maps for the two monkeys. As can be seen, GF and IM neurons had similar locations in the pSTS with the highest density around stereotactic coordinates A0-A2 in monkey T and P1-A1 in monkey L (A0 represents the interaural plane). This hot spot is located on the ventral bank of the pSTS and encroaches on the fundus and the dorsal part of the posterior inferotemporal cortex (pITd). To compare the location of the neurons found in this study with the topography of the GF patch as delineated by a significant contrast between BOLD signals evoked by GF and IM respectively (*6*), we calculated an analogous contrast map based on the electrophysiological heat maps for GF and IM. Despite the general overlap of GF and IM neurons, the contrast map exhibited a clear dominance of GF related activity, due to the larger number of GF neurons. The location of this GF-IM hot spot is comparable to the location of the GFP obtained by BOLD imaging.

**Fig. 5.**
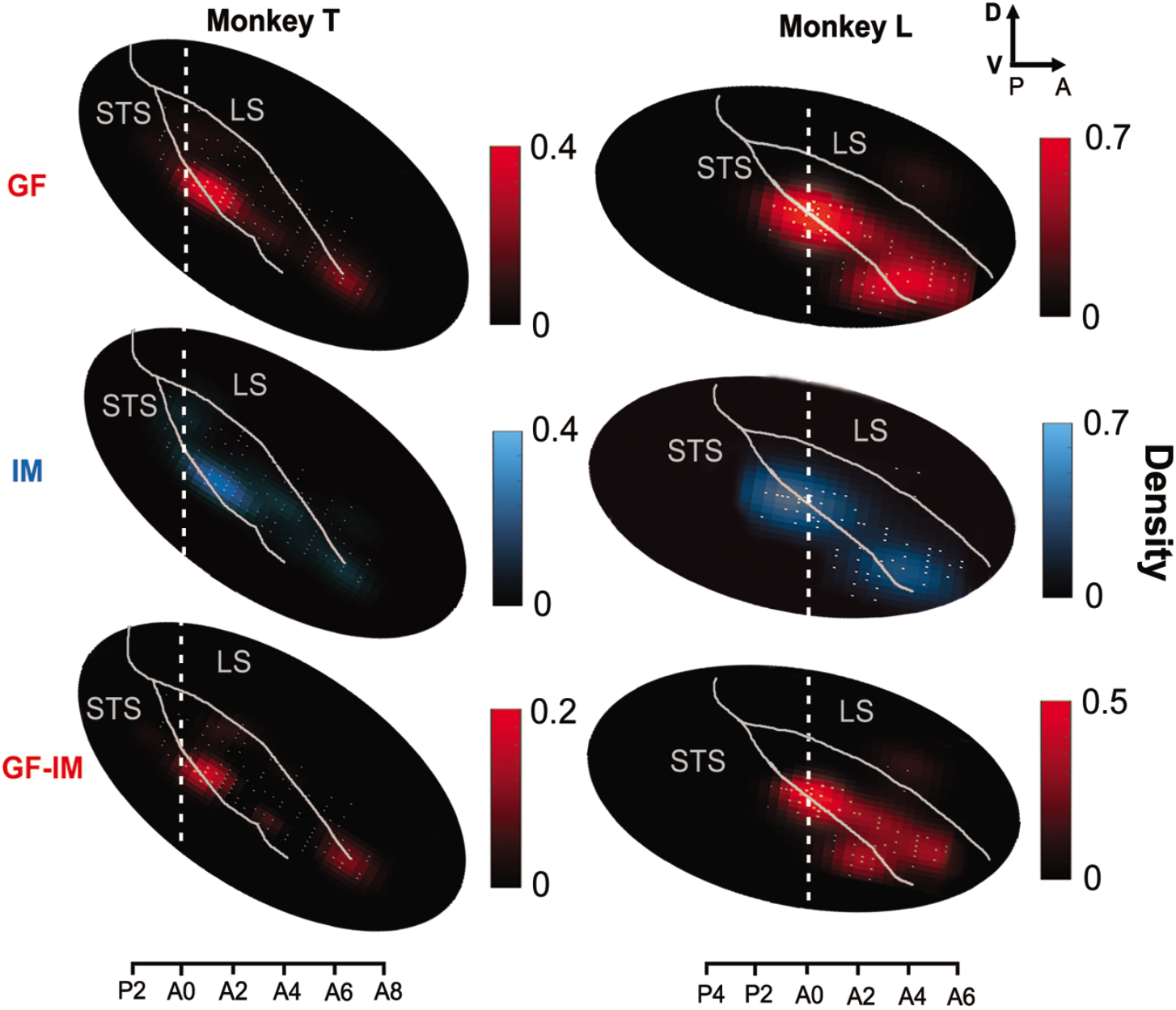
Topography of the GF and IM neurons in the pSTS. Heat maps of the density of GF neurons (red) and IM neurons (blue) in the pSTS of the two monkeys. The two lowest panels depict the contrast between the heat maps of GF neurons and – IM neurons density. A0 is the interaural plane. The density scale represents the number of neurons in elements of 0.5 mm^2^ of the pSTS surface after passing through a 2D spatial filter (Gaussian, σ=2mm).

## Discussion

The posterior STS exhibits a clear functional topography with neurons presenting gaze-following related activity confined to a relatively small area in the lower bank and fundus of the STS around A0-A2 in one monkey and A1-P1 in the other monkey, the gaze-following patch (GFP), clearly separated from neighboring face-selective cortex. The location of the GFP as determined by the properties of single neurons is in good accordance with the location of the GFP as identified by fMRI (*6*). Although boundaries of areas delineated by fMRI are based on somewhat arbitrary statistical thresholds they are often perceived as being sharp and, moreover, associated with qualitatively different functions on both sides. However, our electrophysiological exploration clearly showed that the boundaries of the electrophysiologically defined GFP are gradual with quite a few GF neurons located many millimeters away from the gaze-following hotspot. Shifts of endogenous attention have recently been shown to elicit BOLD activity in the dorsal posterior inferotemporal cortex (pITd) of monkeys (*15*). Considering the published coordinates, this area might be close to the GFP or even overlap with it. Hence, could it be that the GFP is a generic node in an attention network rather than playing a distinct role in gaze-following and joint attention? We think that this possibility can be rejected given the fact that just a few (n=8) of the GF neurons tested responded in a control task in which spatial targets had to be identified by a learned association with distinct abstract objects. On the other hand, some of the spatially-selective GF neurons also exhibited an at least weak interest in shifts of spatial attention guided by learned associations between facial identities and target locations and in general, the regions in which were found GF- and IM related activity overlapped, although the former clearly dominated. In any case, it is the usage of facial information for the purpose of focusing spatial attention that characterizes the GFP. Faces without inherent or learned spatial value do not drive neurons in the GFP. And conversely, the face selective neurons outside the GFP do not respond to shifting spatial attention, no matter whether the faces provide directional information or not. Of course, this does not preclude the possibility, actually suggested by anatomical proximity, that the GFP depends on input from face-selective neurons. The availability of information on spatial targets derived from gaze and in general faces does not necessarily imply a key role in controlling the behavior. Indeed a causal role in guiding behavior is suggested by the fact that the discriminatory power of the population signal on the correct target predicted the behavioral choice. Further support for a causal role comes from a previous study which demonstrated that reversible inactivation of the pSTS compromised the ability of monkeys to use gaze cues to guide target choices (*16*). While the selection of injection sites was based on the response of neurons to a passive face-viewing task and ignorant of physiological landmarks reflecting the preferences of neurons for active GF responses, the reported coordinates suggest that the relatively large injections might have involved the GFP.

By stimulating parietal area LIP, Crapse and Tsao have recently been able to evoke BOLD activity in a region of the monkey pSTS whose coordinates correspond to the GFP (*17*). Hence, it is tempting to speculate that LIP may draw on information on spatial choices prompted by the other’s gaze, originating from the GFP. This input might allow LIP to update its spatial saliency map and to reallocate spatial attention. Such a transfer of information would explain the fact that neurons in LIP present activity related to spatial shifts of attention evoked by gaze cues (*18*).

The GFP does not seem to be confined to ignite gaze-following but also to help suppressing it if not pertinent. This is suggested by the fact that many GFP neurons are sensitive to the rule specifying if the gaze should be followed or not. The observer’s ability to implement a prevailing rule such as to inhibit gaze-following is predicted by the discriminatory power of the population based rule-related activity in a given moment. This suggests a key role of the GFP in controlling gaze-following. The prefrontal cortex is thought to be important for the encoding of rules (*19*). This is for instance indicated by the difficulties of patients with prefrontal lobe damage in following rules (*20*). Hence, it may well be assumed that the rule sensitivity of neurons in the GFP might be a consequence of the integration of top down information from prefrontal cortex. Although we know that output from Brodmann areas 8 and 46 of prefrontal cortex reaches the posterior parts of the STS (*21, 22*), it remains open if the GFP is among the target structures.

Like human gaze-following, also monkey gaze-following seems to be a domain-specific faculty that does not have to be learned from scratch, resorting to domain general machinery. Arguably the latter is needed to learn to associate particular spatial targets with facial identities or abstract objects as required in our study. And we would interpret the existence of identity-mapping related signals in the GFP as reflections of the learned association. Yet, how sure can we be that the gaze-following related activity in the GFP is not also a signature of a learned association? We think that the following arguments render this possibility unlikely. 1. The notion that monkey head gaze-following is domain specific has received substantial support from a previous behavioral study which delineated the position of a monkey’s focus of attention guided by gaze or by identity cues (*13*). Shifts of spatial attention could be fully suppressed if prompted by identity cues. However, shifts of attention guided by gaze cues were blocked only largely, yet not entirely, even after extensive periods of training (*13*). The inability to unlearn gaze-following completely suggests an inborn behavioral capacity not modifiable by learning. 2. A patient suffering from right hemisphere temporal lesion no longer benefitted from gaze direction cues when detecting peripheral targets while the ability to use arrow cues remained intact (*23, 24*). 3. The mark of gaze-following related activity in the GFP is considerably stronger than the one of identity-mapping with the number of GF neurons around 4-fold more (Fig.2a). Hence, the evidence available is in line with the interpretation that the GFP is a central, and possibly domain specific node in a network for the ignition and the control of gaze-following. The emergence of identity-mapping related activity in the GFP is most probably a consequence of the need to control gaze-following based on identity information.

## Acknowledgments

We are grateful to Friedemann Bunjes and Peter W. Dicke for valuable technical support and Ian Chong for critical reading of the manuscript.

## Author contributions

**P. T.** developed the conceptual framework of the research. **P. T.** and **H. R.** designed the experiments, interpreted the results and wrote the paper. **H. R.** performed the experiments and analyzed the data.

## Data and materials availability

Data is available on reasonable request from authors.

## Supplementary Materials

### Materials and Methods

#### Animals, surgery, and recording methods

All experimental preparation and procedures were approved by the local animal care committee (Regierungspräsidium Tübingen, Abteilung Tierschutz) and fully complied with German law and the National Institutes of Health’s *Guide for the Care and Use of Laboratory Animals*. Two male rhesus (Macaca mulatta) monkeys (T and L) of weights 8 kg and 11 kg respectively were used in this study. Before the recording chamber was implanted, we acquired structural Magnetic Resonance Imaging (MRI) scans to identify implant locations. Scans were carried out in a Siemens 3T scanner. Then monkeys were implanted with a titanium head-post to restrain the head during the experiment, scleral search coils for eye position recording and a cylindrical titanium chamber for the introduction of microelectrodes.

Monkey L had participated in a previous fMRI study that had led to the identification of the GFP (*1*). This allowed us to use the stereotactic data available to determine the position and orientation of the chamber on the skull in order to approach the GFP. For the placement of the chamber in monkey T, we relied on the average location of the GFP in the two monkeys that had participated in the fMRI study. All surgeries were carried out under combination anesthesia with isoflurane and remifentanil (1-2 μg/kg/min) with monitoring of all physiological parameters (heart rate, blood oxygen saturation, blood pressure, body temperature). After surgery, opioid analgesics (buprenorphine) were administered until no sign of pain was evident any more. The experiments commenced only after full recovery about 12 days after surgery.

#### Single-unit recording

We recorded single unit activity with vertically movable glass insulated microelectrodes (Alpha Omega, 0.5–1 MΩ at 1 KHz) using conventional techniques. In brief, microelectrodes were driven by a homemade multi-channel micromanipulator attached to the recording chamber in every recording session. Up to four microelectrodes were inserted at the same time with at least 1mm distance to each other. The micromanipulator allowed the selection of microelectrodes positions relative to the chamber walls plane with a spatial resolution of 0.5 mm. Single units were isolated online using the spike waveform matching option of the Alpha Omega SnR system. Quality of isolation was again checked offline and only units whose spikes had been stable throughout the whole session were considered for further analysis.

#### Behavioral tasks

Two monkeys were trained on two “active” tasks requiring either following the head gaze of a demonstrator monkey portrayed on a monitor towards distinct spatial targets or, alternatively, the identification of the same targets based on learned associations with the identity of the portrayed demonstrators. Moreover, they were tested on a “passive” task, requiring fixation of a central target, while a series of behaviorally irrelevant face and non-face images, centred on the target, were presented. In the active tasks, trials started with a white fixation point on a dark background. After 500 ms, a neutral monkey face, centred on the fixation point, always looking straight ahead, appeared. 400ms later, the central fixation changed its color to either red or green, informing the monkey on the rule for target selection to be applied to the upcoming view of an oriented monkey face (“demonstrator”). In case of red, the observer was required to follow the demonstrator’s gaze to one of four spatial targets. The green color required the monkey to make a saccade to the target chosen based on a learned association between the 4 target positions and four possible facial identities, while ignoring gaze orientation. Note that we replaced the green color used to indicate the identity mapping rule by blue in all figures for the sake of better visibility. The demonstrator appeared immediately after the disappearance of the straight face and remained on until the end of the trial. The targets became available together with the onset of the demonstrator. The elimination of the central fixation point 200 or 500 ms after the appearance of the demonstrator served as go-signal for the observer. The monkeys received a drop of water as a reward if they kept fixation of the central fixation point and later made a successful saccade to the target as demanded by the prevailing rule. Trials were aborted if monkeys were not keeping their eyes within a window of 2° around the fixation point and the target respectively and were unable to reach the target within 300ms after the go-signal.

The images of straight and oriented faces had a size of 5.6° x 5.6° and were presented in the center of a monitor placed at a distance of 60 cm from the observer. Spatial targets were small red dots (diameter of 0.8°) and were aligned on a virtual horizontal line 1° below the center of the portraits at horizontal eccentricities of −10°, −5°, 5° and 10° with respect to the observer monkey (−40°, −20°, 20° and 40° with respect to the demonstrator monkeys). As the portrait of each individual monkey could be shown in four different head gaze orientations, corresponding to the four spatial targets, the stimulus set involved 16 stimuli (**Fig. 1B**). We used an open source recording and stimulation system for recording of eye movement data and presenting the stimulus images (<u>nrec.neurologie.uni-tuebingen.de/nrec</u>). Gaze-following and identity mapping trials were identical in visual terms except for the color of the instruction cue, available in a short period only, and identical with respect to the motor responses required. Hence, any differences in the associated neuronal responses outside the short presence of the instruction cue had to be a consequence of differences in cognitive strategies and operations.

Finally, the monkeys had to perform the aforementioned passive viewing task in which images of faces and non-face stimuli, centered on the fixation dot, were presented and the monkeys had to keep fixation of the central fixation point (**Fig. 1C**). In this task, we used the same set of images used in a previous study (*1*) in addition to the 16 monkey portraits used in the active tasks and additional 16 human faces (4 identities with four gaze direction similar to those monkeys head direction in the active tasks taken from the Radboud Face Database (*2*)). Monkeys saw in total 144 images of 6°x6°, each lasting for 400 ms and followed by a 400 ms black and white random dot background (pixel size 0.05°). Monkeys were rewarded in this experiment if they were keeping their eyes within a window of 2°x2° around the central fixation point for each image.

#### Statistical Analysis

In order to characterize the discharge patterns evoked in the two active tasks, we determined the mean discharge rate in 3 periods: (*1*) the baseline period: the last 100 ms of the portrait fixation period right before the onset of the rule presentation period, (*2*) the rule period: the 400 ms after onset of the instructive cue, and (*3*) the spatial information period: the period during which the demonstrator was available and the observer waited for the go signal cue. The duration of this latter period was either 200 or 500ms. We refrained from considering later periods because we expected them to be influenced by a complicated mixture of variables like saccade execution, saccade induced visual stimulation, reward expectancy and preparation or outcome evaluation. We determined the task related preferences of neurons by comparing the mean firing rates in the three periods by a non-parametric 1-way ANOVA (Kruskal-Wallis test) (considering p<0.05). When an effect in the 1-way ANOVA was found, the specific phase (rule or spatial cue periods) significantly different from baseline was identified by means of a post-hoc analysis (p < 0.05 Bonferroni corrected for multiple comparisons). Neurons which exhibited a significant change of their discharge rate in the rule and/ or the spatial cueing period were selected for further analysis. A neuron was considered to be spatially selective in the spatial cueing period if its firing rates (only correct trials considered) to the four different targets were significantly different (Kruskal–Wallis one-factor ANOVA, P < 0.05, carried out separately for gaze-following and identity-mapping). The population responses associated with the most and the least preferred target were compared by a Mann-Whitney U test (p<0.05).

A rule selectivity index (RSI), with a theoretical maximum of 1 for gaze following, −1 for identity mapping and a theoretical minimum of 0 for unselective neurons, was calculated for the 400 ms rule according to

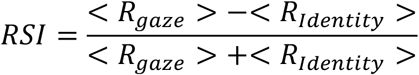

In a similar way, a spatial selectivity index (SSI), with a theoretical maximum of 1 and minimum of 0, was calculated for the 200/500 ms spatial cueing periods according to

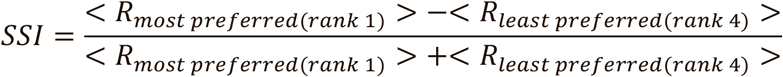

with the operator < > denoting the mean firing rate in correct trials. Changes in the distribution of the RSI and SSI values were evaluated by means of Mann-Whitney U tests.

In the decoding analysis, we deployed a support vector machine (SVM) with 5-fold crossvalidation to determine how well the population discharge predicted the spatial target selected and the choice of the monkeys, considering the prevailing rule. We obtained a decoding time course by performing this analysis in a100 ms window, advanced in steps of 20 ms throughout the whole spatial cueing period. We performed this analysis separately for the two different spatial cueing periods of 200 ms and 500 ms when testing for target location sensitivity. Error trials were classified according to the class of correct trials that they resembled most. Standard errors of the decoding performance were obtained by bootstrapping (n=1000).

### Supplementary Text

#### Low spatial selectivity of face-selective neurons

We also recorded the activity of the same pSTS neurons tested in the two active tasks during passive vision of face and non-face stimuli in order to investigate to what extent the spatially selective neurons are the same classically face-selective neurons. 23 out of 334 neurons tested exhibited significantly stronger responses to the passive vision of faces than to non-face stimuli (p<0.05; Wilcoxon signed-rank test comparison of mean response to faces and non-face stimuli). Out of this total of 23 face-selective neurons sampled from both monkeys, only 6 were from the set of 109 gaze following neurons with spatial selectivity. **Fig. S1** compares the population response of these face-selective neurons in the three tasks, as well as the amount of their spatial modulation for correct and incorrect trials. As can be seen, the average SSI values were very similar and statistically not different for correct and incorrect trials. Also, the response profiles for the most preferred and least preferred targets did not show any significant difference in the spatial cueing period (Mann Whitney U test, p>0.05). Hence, these neurons cannot have immediate relevance for the behavioral choices.

#### Abstract symbol-matching responses

We also investigated if neurons activated in the identity mapping task might also be responsive to learned associations between non face images and spatial locations. To this end, we trained the monkeys to associate four abstract symbols (square, circle, triangle, star) almost the same size (5° x 5°) with the four targets. The trial structure and timing were similar for the GF and IM tasks with the exception that there was no change in the color of the central fixation which could serve as a rule and the four abstract symbols were presented right after the same portrait fixation face used in the previous tasks. As this control task was usually run at the end of a session, once sufficient data in the three main tasks had been collected, only a minority of neurons could be tested on this abstract symbol mapping control. In many other cases, the monkeys were no longer motivated to work or the quality of the spike isolation had deteriorated too much. We only considered neurons for which we had collected at least 8 correct trials for each of the four spatial targets. Both monkeys learned the task well and their performance was very similar to their performance in the gaze-following and identity-mapping tasks (monkey T: mean ± std= 73 ± 1%, monkey L: mean ± std= 74 ± 2% correct). In total, we could test 131 out of the 426 neurons tested on the gaze following and the facial identity mapping task (44 neurons from monkey L and 87 from monkey T) in the symbol mapping task. Only 8 out of the 131 neurons (2 from monkey L, 6 neurons from monkey T) exhibited spatial selectivity, characterized by significant target specific responses in the spatial cueing period (1-way ANOVA (Kruskal-Wallis test, p<0.05). In 3 of the 8 neurons, the number of error trials was sufficiently large to allow a comparison of the responses between correct and incorrect trials. These comparisons did not reveal any significant difference (Mann Whitney U test, p>0.05). Hence, rather than reflecting spatial selectivity, these may be more elementary visual responses evoked by particular abstract objects. On the other hands, given the low probability of neurons exhibiting significant responses to the presence of abstract symbols (8 out of 131 neurons tested; i.e. 6% of the population), these neurons may simply be statistical artifacts.

**Fig. S1.**
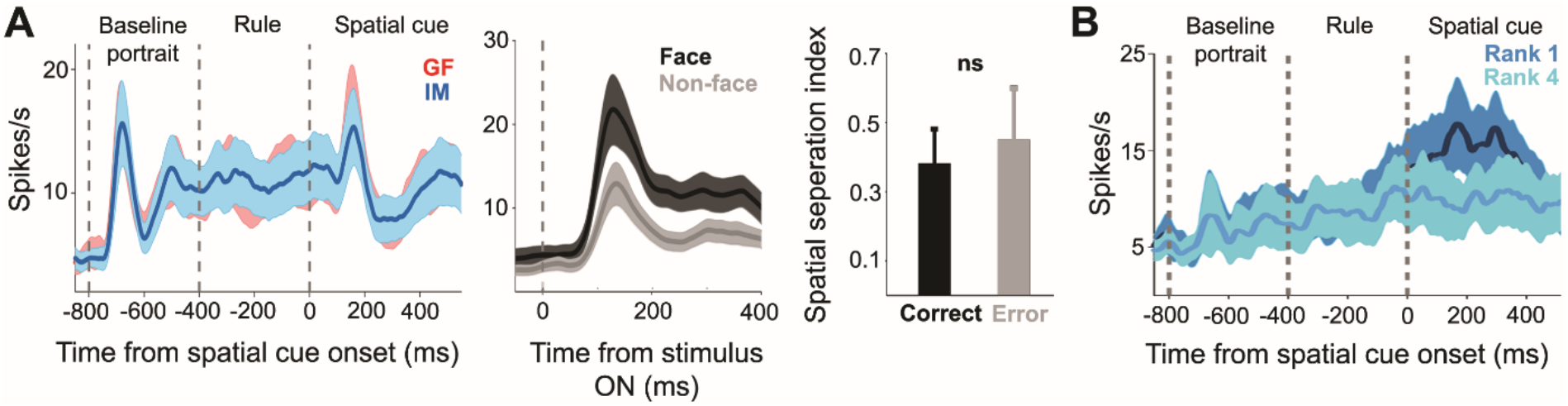
Population response profile of the passive face-selective and IM neurons. **(A)** The population discharge profiles of classical face-selective neurons, exhibiting significantly larger response to faces as compared to non-face stimuli when tested in the passive viewing task. Both profiles show significant responses to the presence of the portraits in both the baseline portrait and the spatial cueing periods, yet, without any difference between the gaze following and the identity matching tasks. Moreover, the population discharge did not differentiate between the most and the least preferred targets, no matter if spatial choices were correct or not. Hence, they hardly contribute to shaping monkeys’ spatial choices. **(B)** The population discharge profile of the spatially selective IM neurons did not differentiate significantly between the most preferred and the least preferred IM targets. Error bars and shaded areas represent standard error in all subfigures.

### Movie S1

Exemplary neuron exhibiting a burst of activity for shifts of the experimenter’s gaze to the left.

